# NK Cells Effectively Mediating Antibody-Mediated Kidney Allograft Rejection Requires a Specific Activation Receptor and Graft Expression of the Ligand

**DOI:** 10.64898/2026.03.03.709363

**Authors:** Yuki Maruyama, Daigo Okada, Hidetoshi Tsuda, Danielle D. Kish, Karen S. Keslar, Nina Dvorina, William M. Baldwin, Robert L. Fairchild

## Abstract

Acute antibody-mediated rejection (aABMR) is an important cause of clinical kidney graft injury and failure. Transcripts associated with NK cell activation in graft biopsies are diagnostic of aABMR, but mechanisms underlying NK cell activation during ABMR remain poorly understood. In contrast to the long-term (> 60 days) survival of complete MHC-mismatched kidney allografts in wild type C57BL/6 mice, B6.CCR5^⁻/⁻^ recipients develop high titers of donor-specific antibody (DSA) with allograft rejection between days 18 to 28 post-transplant. This has allowed investigation of mechanisms underlying NK cell activation within kidney allografts during aABMR. DSA titers first became detectable in B6.CCR5^⁻/⁻^ (H-2^b^) recipients of A/J (H-2^a^) kidney allografts at day 8 and peaked on day 15 post-transplant and was accompanied by a parallel increase in mRNA levels of Rae-1e, a ligand for the NK cell activation receptor NKG2D. A/J kidneys in B6.CCR5^⁻/⁻^NKG2D^⁻/⁻^ recipients and A/J.Rae-1e^⁻/⁻^ kidneys in B6.CCR5^⁻/⁻^ recipients survived >60 days, despite high serum DSA levels. Flow cytometric analysis of allograft infiltrating cells in B6.CCR5^⁻/⁻^ recipients on day 15 post-transplant revealed inflammatory monocyte and NK cell infiltration and NK cell activation to proliferate and express CD107a, a marker of cytotoxic function. These features of aABMR were absent or markedly reduced by recipient NKG2D- or donor graft Rae-1e-deficiency. These findings suggest that interference with expression of allograft Rae-1e or recipient NK cell NKG2D abrogates aABMR despite persistently high DSA levels and that aABMR requires coordination between infiltrating NK cell and inflammatory monocyte activation within the kidney allograft.

## Introduction

Antibody mediated rejection causes substantial acute and chronic graft injury and continues to be an important problem undermining the success of clinical kidney transplants (1, 2). While effective strategies have been developed to prevent or treat ongoing T cell mediated rejection (TCMR), there remains a lack of effective therapies to arrest ongoing ABMR and prevent its reoccurrence. Sensitization to donor allogeneic MHC molecules generates antibodies that bind to the class I and/or class II MHC molecules expressed on the graft endothelium. These donor specific antibodies (DSA) mediate direct endothelial cell injury through activation of complement as well as the recruitment of myeloid cells and other innate immune cells to exacerbate inflammation and graft cell injury (3–5). In addition to the detection of serum DSA, the margination of monocytes, macrophages and neutrophils in peritubular capillaries is a diagnostic histopathologic feature of ongoing ABMR. Another important component of innate immune responses during ABMR is NK cell activation in the graft microvasculature. Studies by Hidalgo and colleagues (6, 7) reported kidney allograft transcript profiles expressed during ABMR, but not TCMR, include those associated with NK cell activation and are used diagnostically to assess ABMR While the presence of NK cell responses during ABMR is recognized, how the activation of NK cells and other innate immune components is organized to mediate the graft vascular injury during ABMR remains unclear and underlies the need for models that include cellular and molecular features of the injury observed in clinical kidney transplants.

Whereas complete MHC mismatched kidney grafts survive long-term in wild type C57BL/6 mice, the allografts are rejected within 18-28 days after transplant in B6.CCR5^-/-^recipients (8–12). This rejection requires the high titers of DSA produced by the CCR5-deficient recipient and shares histopathologic features with ABMR of clinical kidney transplants including, C4d deposition in the capillaries as an indicator of DSA binding to target allogeneic MHC targets, margination of myeloid cell and neutrophils in peritubular capillaries, and fibrin deposition in the glomeruli. Results from this model have also directly shown NK cell activation within the kidney allografts and that either antibody-mediated depletion of NK cells or inhibition of their activation abrogates aABMR (10–13), resulting in prolonged graft survival beyond day 60 post-transplant despite the continued presence of high serum DSA titers. The presence or absence of NK cell activation within kidney allografts in this model is further indicated by graft tissue expression of the NK cell associated transcripts detected in clinical kidney graft biopsies during ABMR (14, 15).

NK cell activation within inflammatory vascular sites requires a skewing of the balance of positive and negative signals from two sets of receptors (16, 17). NK cell activation is negatively regulated by receptors that engage self-class I MHC molecules, so this missing inhibitory self-signaling is absent during responses to complete MHC mismatched kidney allografts. Positive signaling for expression of NK cell effector function, including IFN-γ and granzyme B, is delivered through activation receptors that engage ligands expressed on target cells on cells in the allograft, including stress induced ligands such as heat shock proteins and Rae-1e (18, 19). Whether expression of these ligands is generated at the time ischemic kidney grafts are reperfused during transplantation or are induced by the binding of DSA to target donor MHC on the vascular endothelium remains unclear. Furthermore, NK cells express multiple activation receptors suggesting conceivably blocking only one receptor may not effectively inhibit NK cell activation (17, 19). The NK cell-dependent ABMR of kidney allografts in CCR5^-/-^ recipients afforded the opportunity to test the role of such receptors and their ligands in NK cell activation within the allografts and gain further insights into mechanisms underlying aABMR in this model.

## Results

### Expression of Stress Ligands during ABMR

To gain insights into components required for activation of NK cells during ABMR of A/J kidney allografts in B6.CCR5^-/-^ recipients, kidney iso- and allo-grafts were harvested on day 14 post-transplant when DSA titers and NK cell activation within the allografts reach peak levels. Whole graft RNA was isolated and tested by qPCR for expression of transcripts encoding a panel of potential ligands for NK cell receptors (Figure 1A). Whereas Rae-1e, a ligand for NKG2D, Rae-1d, and H60b were expressed in allo- but not iso-grafts at this time point, transcripts encoding H60a, IL-33 and ST2 were at either lower or absent levels in allo- vs. iso-grafts. Temporal analysis of expression indicated that Rae-1e and H60b transcripts were evident at low levels in the graft on day 10 post-transplant and continued to rise on day 15 commensurate with the detection and rise of serum DSA in the serum of the allograft recipients (Figure 1B and 1C).

**Figure 1.**
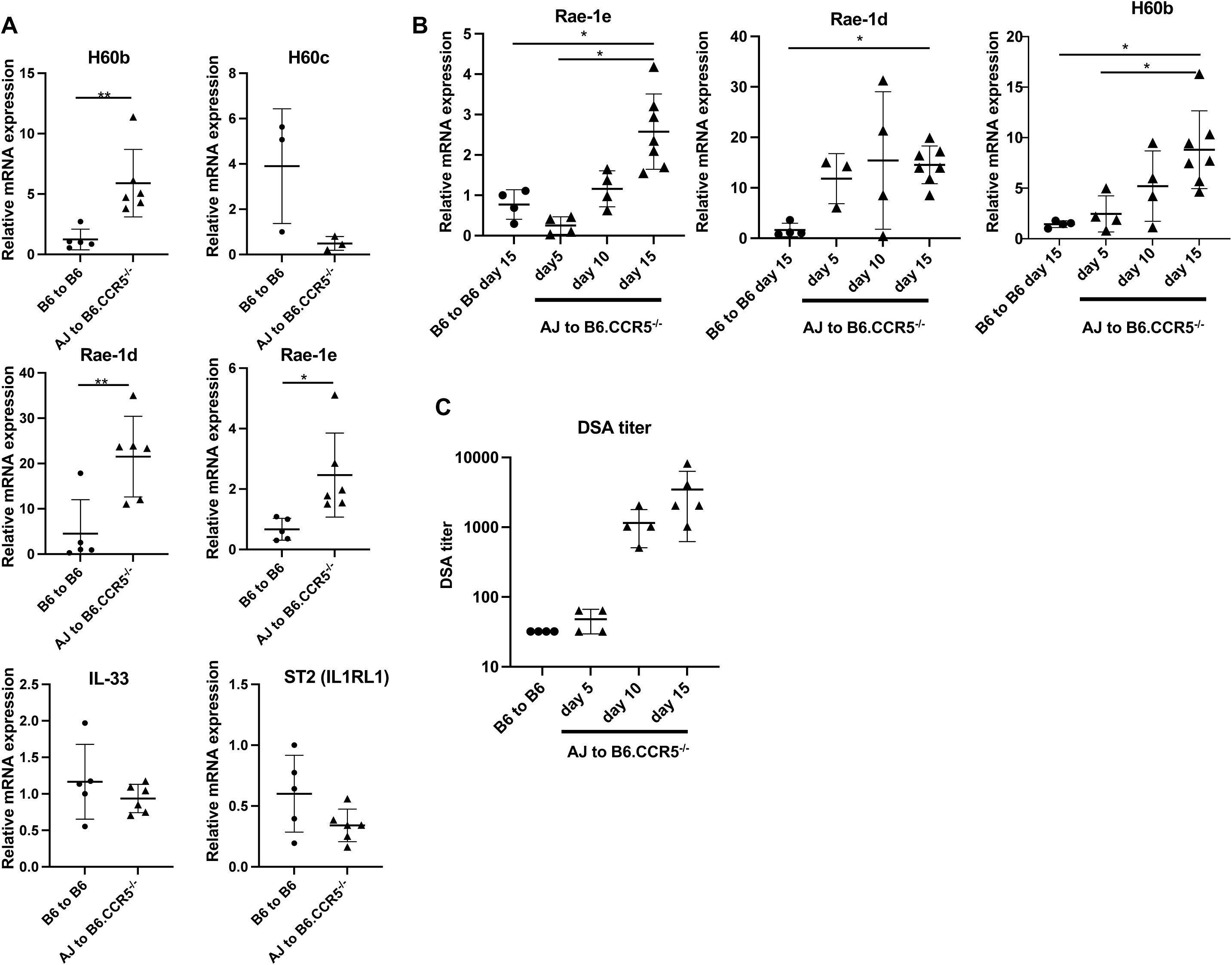
Determination of ligands for NK cell activation receptors during ABMR in kidney allografts in CCR5^-/-^ recipients. (A) RNA was isolated from homogenates of groups iso – (B6 to B6) and allo- (A/J to B6.CCR5^-/-^) graft recipients on post-operative day (POD) 15 post-transplant. Quantitative PCR analysis of H60b, H60c, Rae-1d, and Rae-1e, as well as IL-33 and ST2 (IL1RL1) and relative mRNA expression levels in AJ grafts to B6.CCR5^⁻/⁻^ recipients (n=6) are shown in comparison with a B6 graft to B6 recipient (n=4). Data indicate the mean ± SD expression levels for each group. *P < 0.05, **P < 0.01 as determined by 2-tailed t test. (B) qPCR analysis of Rae-1e, Rae-1d, and H60b expression in AJ grafts to B6.CCR5^⁻/⁻^ recipients at POD 5 (n=4), 10 (n=4), and 15 (n=7), compared with B6 grafts to B6 recipients at POD15 (n=4). Data indicate the mean ± SD. *P < 0.05 as determined by one-way ANOVA test. (C) Serum samples were obtained from individual B6 recipients of B6 grafts at POD15 (n=4) and from B6.CCR5^⁻/⁻^ recipients of AJ grafts at POD5 (n=4), 10 (n=4), and 15 (n=5) and donor-specific antibody (DSA) titers were determined. Data are presented as the mean DSA titer ± SD for each transplant recipient group.

### Impact of Rae-1e Allograft and Recipient NKG2D Deficiencies on aABMR

The role of Rae-1e was further investigated as an allograft ligand activating NK cells to mediate acute kidney graft injury during ABMR. Rae-1e^-/-^ mice on the C57BL/6 (H-2^b^) were backcrossed 10 generations to the kidney allograft donor A/J (H-2^a^) mouse background to generate A/J.Rae-1e^-/-^ mice and were transplanted to groups of B6.CCR5^-/-^ recipients and survival compared with kidney allografts from wild type A/J donors (Figure 2A). Whereas wild type A/J kidney allografts rejected between days 20 and 30, all A/J.Rae-1e^-/-^ kidneys survived beyond day 58 post-transplant (Figure 2B), despite the sustained presence of high serum DSA titers (Figure 2C). Consistent with this prolonged survival, all wild-type A/J kidney allografts survived more than 60 days after transplant to B6.CCR5^-/-^NKG2D^-/-^ recipients even though the titers of DSA generated were similar in B6.CCR5^-/-^ and B6.CCR5^-/-^NKG2D^-/-^ recipients of A/J kidney allografts (Figure 2C and 2D). Similarly, treatment of B6.CCR5^-/-^ recipients of A/J kidney allografts with anti-NKG2D mAb beginning at the time serum DSA was first detected on day 8 post-transplant extended allograft survival in most treated recipients past day 60 post-transplant, also without affecting the DSA titers (Figure 2C and 2D). Flow cytometry analyses indicated that treatment with anti-NKG2D mAb depleted NK1.1^+^DX5^+^NKG2D^+^ cell populations in the spleen and in the allograft without affecting the presence of the NK1.1^+^DX5^+^NKG2D^+^ cells in each site (data not shown).

**Figure 2.**
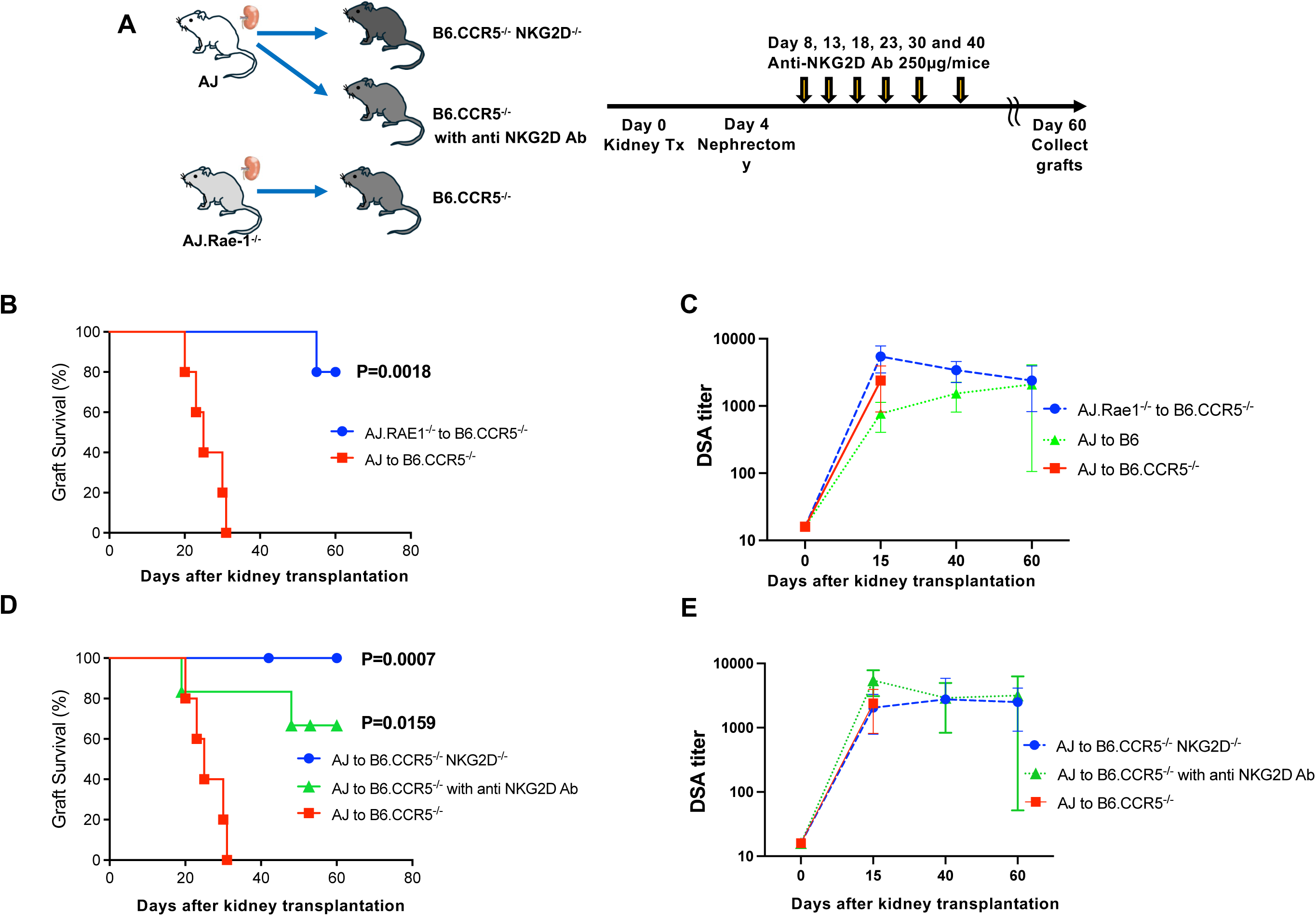
Rae-1 deficiency in allografts and recipient NKG2D deficiency abrogates ABMR and prolongs kidney allograft survival. (A) Experimental design and protocol: AJ or AJ.Rae-1^⁻/⁻^ donor kidneys were transplanted into B6.CCR5^⁻/⁻^ recipients, B6.CCR5^⁻/⁻^NKG2D^⁻/⁻^ recipients, or B6.CCR5^⁻/⁻^ recipients treated with anti-NKG2D mAb (250 µg/mouse administered on days 8, 13, 18, 23, 30, and 40 after transplantation). Nephrectomy of the native kidney was performed on day 4. (B) AJ (n=6) or AJ.Rae-1^⁻/⁻^ (n=5) kidney allografts were transplanted to groups of B6.CCR5^-/-^ recipients and allograft survival was monitored. All AJ.Rae-1e^⁻/⁻^ kidneys survived for more than 55 days after transplantation with significantly prolonged graft survival compared with A/J kidney allografts by Kaplan–Meier analysis with log-rank test (p = 0.0018). (C) Serum from wild-type C57BL/6 or B6.CCR5^-/-^ recipients of A/J or AJ.Rae-1^⁻/⁻^ kidney grafts was obtained from individual recipients at the indicated times after transplant and the DSA titers. Data indicate mean DSA titer for each graft recipient group ± SD. (D) AJ kidney allografts were transplanted to groups (n = 6/group) of B6.CCR5^-/-^ recipients and allograft survival was monitored. Kaplan–Meier analysis with log-rank test was used to analyze graft survival in B6.CCR5^⁻/⁻^, B6.CCR5^⁻/⁻^NKG2D^⁻/⁻^, and B6.CCR5^⁻/⁻^ recipients treated with anti-NKG2D mAb transplanted AJ kidney allografts. Both genetic deletion of NKG2D and antibody-mediated blockade significantly prolonged graft survival compared with B6.CCR5^⁻/⁻^ controls (p=0.0007 and p=0.0159, respectively). (E) Serum DSA titers in B6.CCR5^⁻/⁻^, B6.CCR5^⁻/⁻^NKG2D^⁻/⁻^and B6.CCR5⁻^/^⁻ treated with anti-NKG2D mAb transplanted AJ kidney allografts., and AJ→B6.CCR5⁻^/^⁻ recipients at the indicated time points. Data indicate mean titer for each graft recipient group ± SD

Histolopathologic evaluation of allografts from each group was performed on day 15 post-transplant. All allografts, including wild type A/J kidneys transplanted to wild type C57BL/6 recipients, exhibited diffuse C4d deposition in peritubular and glomerular capillaries consistent with the engagement of DSA to their target MHC molecules in the allografts (Figure 3A). In addition, dilation of peritubular capillaries was clearly evident in A/J allografts from B6.CCR5^-/-^recipients, but not in the non-rejecting allografts from wild type C57BL/6 and B6.CCR5^-/-^NKG2D^-/-^ recipients and in A/J.Rae-1e^-/-^ allografts from B6.CCR5^-/-^ recipients. The capillary dilation in the rejecting A/J allografts from B6.CCR5^-/-^ recipients was accompanied by severe cortical edema that was not observed in allografts from the other recipient groups (Figure 3B and data not shown).

**Figure 3.**
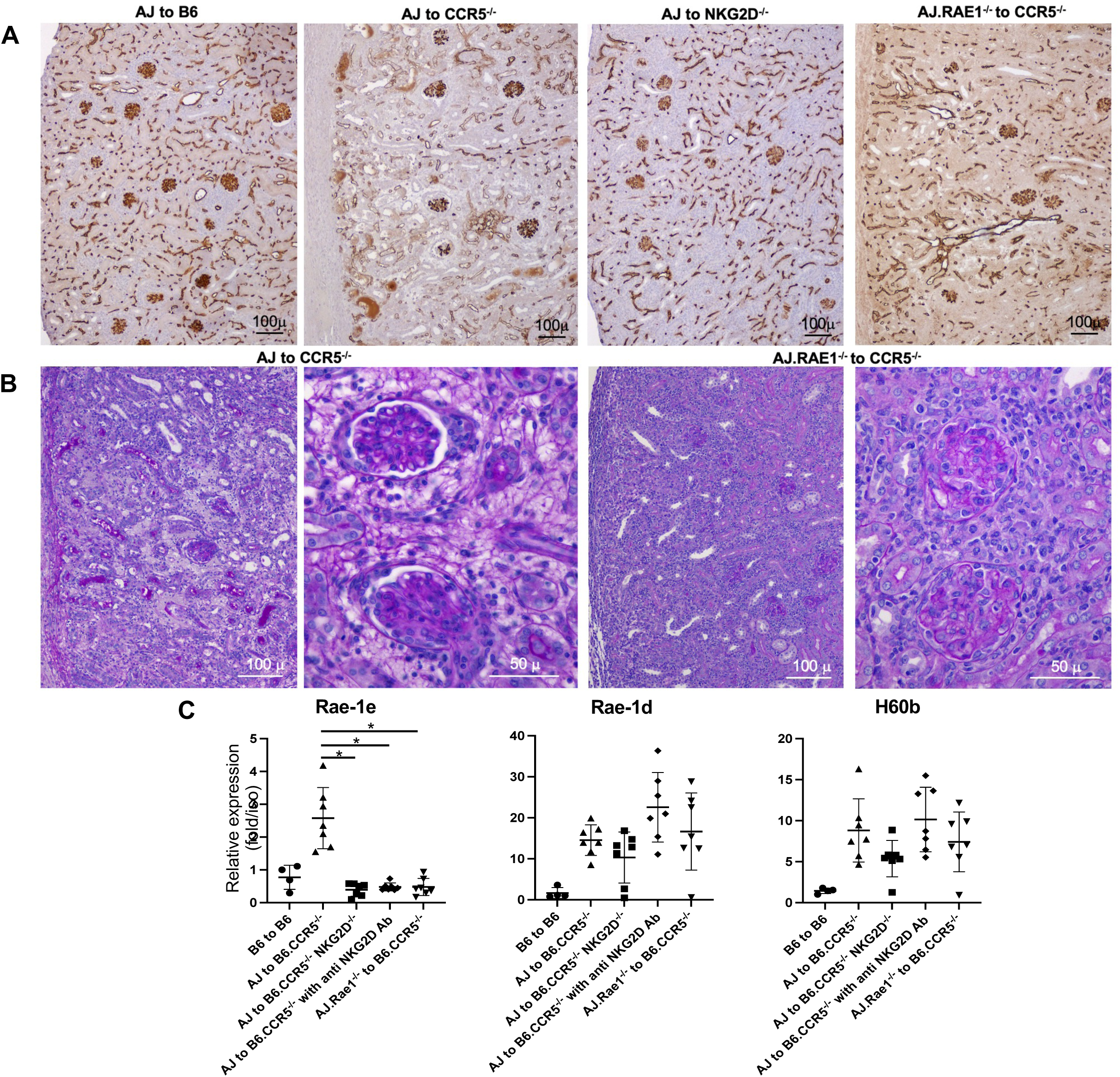
Suppression of NKG2D signaling by Rae-1 deficiency mitigates graft injury independently of complement deposition. (A) Representative C4d immunohistochemical staining of kidney allografts at day 15 POD. Immunohistology demonstrates diffuse strong deposition of C4D in peritubular and glomerular capillaries of AJ kidney allografts to wild-type C57BL/6, B6.CCR5^⁻/⁻^, B6.CCR5^⁻/⁻^NKG2D^⁻/⁻^ and AJ.Rae-1^⁻/⁻^ kidney allografts to B6.CCR5^⁻/⁻^ recipients with capillary dilation observed in A/J to B6.CCR5^⁻/⁻^ recipients. (B) Periodic acid–Schiff (PAS) staining of kidney allografts reveals cortical edema and dilation of peritubular capillaries in AJ graft to B6.CCR5^⁻/⁻^. In contrast, AJ.Rae-1^⁻/⁻^ grafts to B6.CCR5^⁻/⁻^ showed attenuated tissue injury with reduced edema. Scale bars, 100 μm (low power) and 50 μm (high power). (C) Quantitative PCR analysis of NKG2D ligands (Rae-1e, Rae-1d, and H60b) in kidney grafts on day 15 POD. Relative mRNA expression levels in each group are shown in comparison with a B6 graft to B6 recipient. Rae-1e expression was significantly reduced in AJ grafts to B6.CCR5^⁻/⁻^NKG2D^⁻/⁻^ and B6.CCR5⁻^/^⁻ treated with anti-NKG2D mAb and AJ.Rae-1^⁻/⁻^ grafts to B6.CCR5^⁻/⁻^ recipients compared with expression in AJ grafts to B6.CCR5⁻^/^⁻ recipients; whereas, Rae-1d and H60b expression showed no significant differences in allograft groups. Data indicate the mean ± SD. *P < 0.05 as determined by one-way ANOVA test.

The expression of transcripts encoding Rae-1d, Rae-1e, and H60b in kidney grafts was also determined on day 15 post-transplant for each of the recipient groups (Figure 3C). Rae-1e transcripts were significantly expressed at high levels in the rejecting A/J allografts from B6.CCR5^-/-^ recipients vs. low-absent expression in the non-rejecting grafts from the other recipient groups. In contrast, both Rae-1d and H60b were expressed in allografts from all of the recipient groups at this time point post-transplant.

Transcripts encoding other mediators of ABMR were also measured in the kidney grafts on day 15 post-transplant and compared between the recipient groups (Figure 4). Similar to Rae-1e, expression of effector function transcripts (IFN-γ, granzyme B, perforin, and TNFα) were significantly increased in the rejecting A/J allografts from B6.CCR5^-/-^ recipients vs. the expression in grafts from the other recipient groups. This significantly elevated expression pattern in A/J kidney allografts from B6.CCR5^-/-^ recipients vs. grafts from the other groups was also observed for transcripts encoding the monocyte chemoattractant CCL2, the NK cell chemoattractants CXCL9 and CXCL10, and for CX_3_CR1, which is expressed on NK cells and myeloid cells and directs their recruitment to inflammatory sites. Interestingly, the ABMR diagnostic transcripts MyBL1 and Sh2D1B1 were expressed in allografts, not isografts, from all of the recipient groups producing DSA and with or without rejection of their grafts.

**Figure 4.**
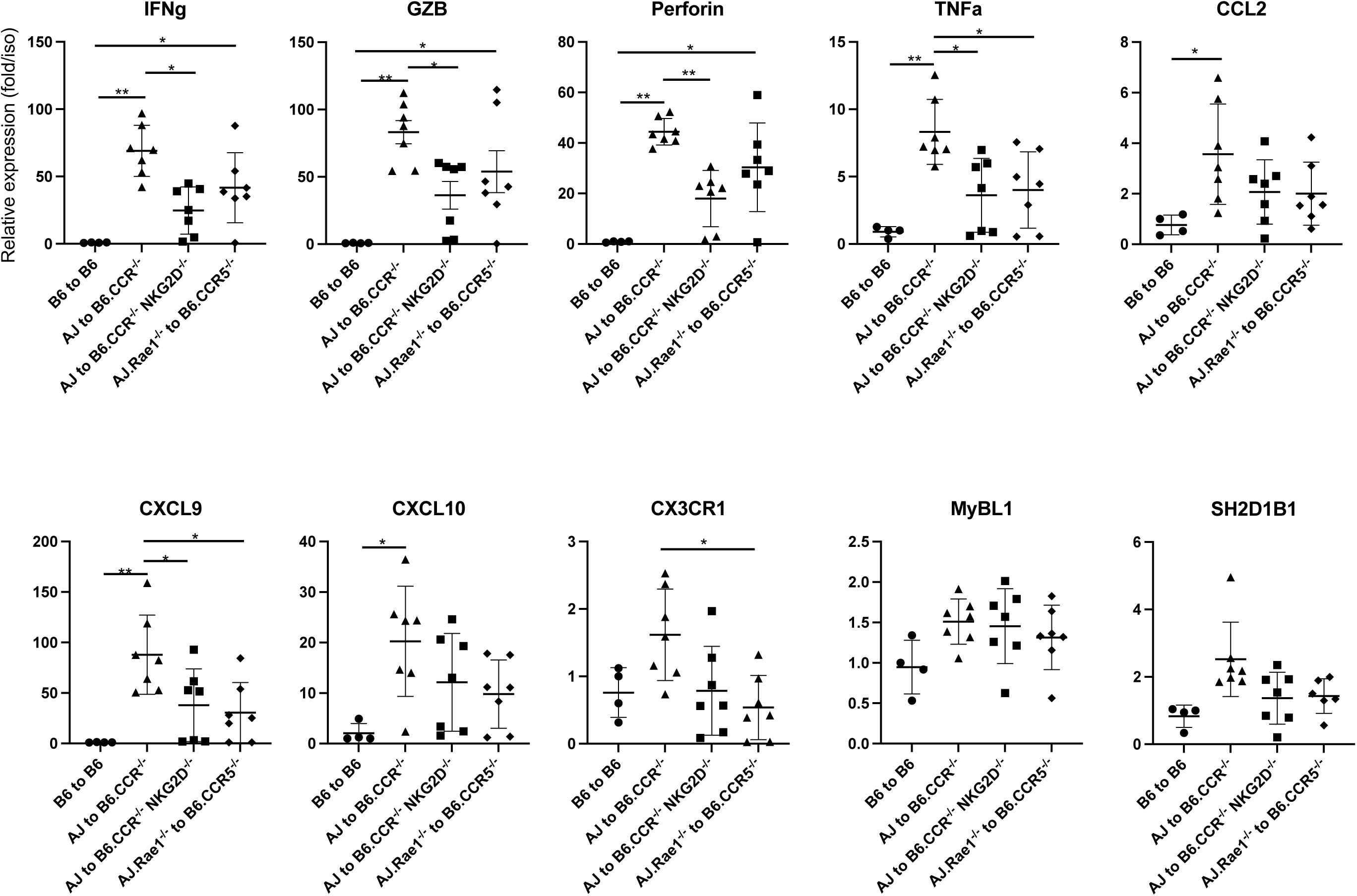
NKG2D signaling regulates the expression of ABMR-associated inflammatory mediators in kidney allografts. Quantitative PCR analysis of transcripts encoding mediators associated with antibody-mediated rejection (ABMR) was performed in kidney grafts harvested on day 15 POD. The expression levels of IFN-γ, Gzmb (GZB), Perforin, TNFα, CCL2, CXCL9, CXCL10, CX3CR1, MyBL1, and SH2D1B1 were compared between B6 isografts to B6 recipients, AJ allografts to B6.CCR5^⁻/⁻^, A/J allografts to B6.CCR5^⁻/⁻^NKG2D^⁻/⁻^, and AJ.Rae-1^⁻/⁻^ grafts to B6.CCR5^⁻/⁻^ recipients. Relative mRNA expression levels in each group are shown in comparison with a B6 graft to B6 recipient. Data are presented as mean ± SD. Statistical significance was determined by one-way ANOVA with appropriate post hoc comparisons. Data indicate the mean ± SD. *P < 0.05, **P < 0.01 as determined by one-way ANOVA test.

### Absent NK cell activation in Rae-1 deficient allografts and wild type allografts in NKG2D deficient recipients

The activation of NK cells within the allografts on day 15 was directly assessed in each of the 4 recipient groups. In contrast to the number of NK cells accumulating in wild type A/J kidney allografts on day 15 post-transplant, there was a significant 50-75% decrease in NK cell numbers within A/J.Rae-1e^-/-^ allografts harvested from B6.CCR5^-/-^ recipients and in A/J allografts harvested from B6.CCR5^-/-^NKG2D^-/-^ recipients (Figure 5A and B). These decreases in NK cell numbers in A/J kidney allografts were also observed when B6.CCR5^-/-^ recipients were treated with anti-NKG2D mAb (Figure 5A and B).

**Figure 5.**
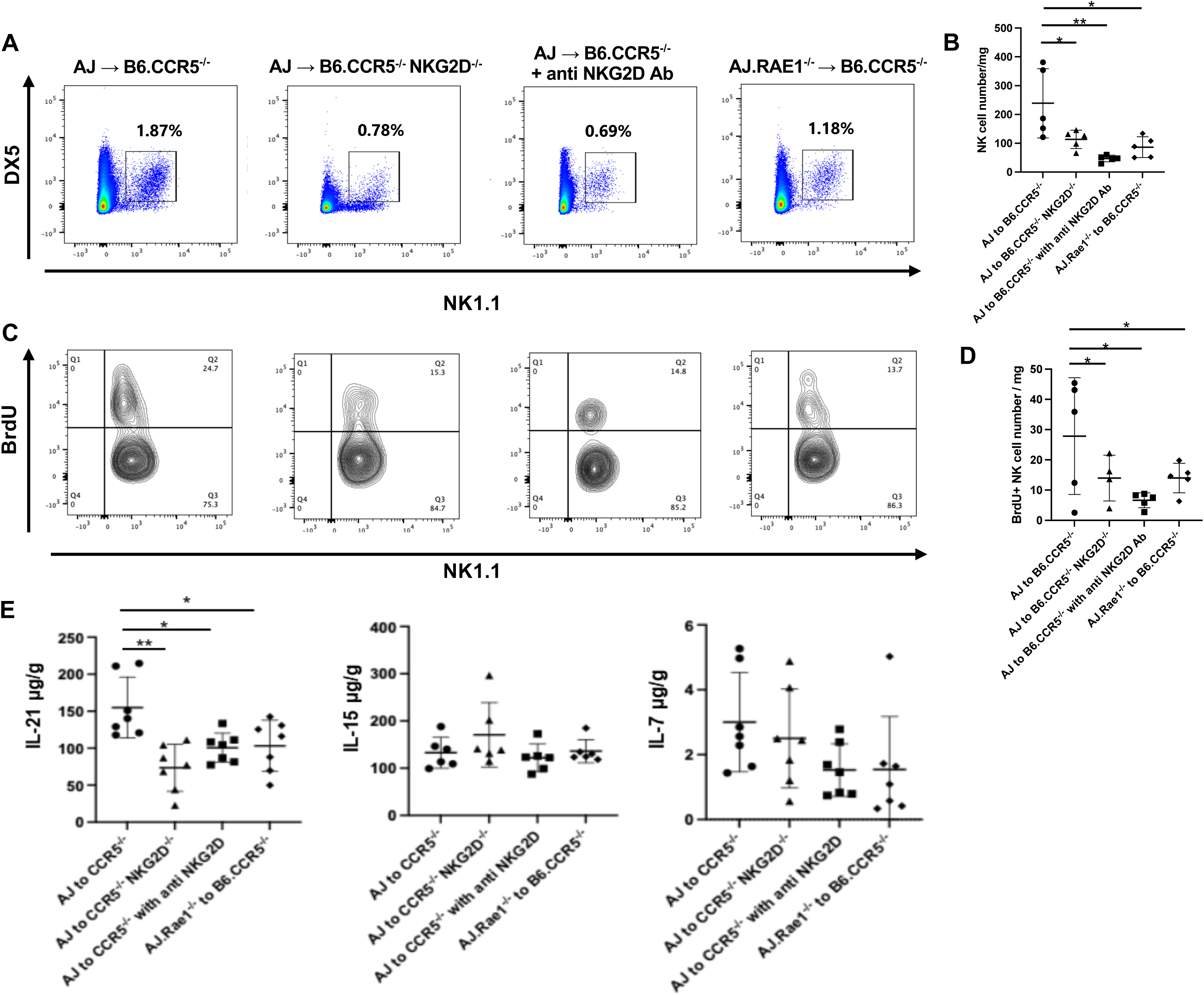
Absence of recipient NKG2D or donor Rae-1 suppresses NK cell accumulation and proliferation in kidney allografts. AJ kidney allografts were transplanted to groups (n = 5/group) of B6.CCR5^-/-^, B6.CCR5^⁻/⁻^NKG2D^⁻/⁻^, B6.CCR5^⁻/⁻^ treated with anti-NKG2D mAb recipients and in AJ.Rae-1^⁻/⁻^ allografts transplanted to B6.CCR5^⁻/⁻^recipients. (A) Representative flow cytometric analysis of infiltrating NK cells in kidney allografts on day 15 post-transplant. Single-cell suspensions were prepared from graft tissues and gated on CD45⁺ leukocytes, followed by exclusion of CD3⁺ cells. NK cells were defined as NK1.1⁺DX5⁺ cells within the CD45⁺CD3⁻ population. (B) Quantification of total NK cell numbers per mg of graft tissue in the indicated groups. Absence of recipient NKG2D or donor Rae-1 suppresses NK cell accumulation. Data indicate the mean ± SD. *P < 0.05, **P < 0.01 as determined by one-way ANOVA test. (C) Representative flow cytometric analysis of BrdU incorporation in graft-infiltrating NK cells. BrdU was administered i.p. on POD 14 and allografts were harvested and cell aliquots stained to detect BrdU^+^NK1.1^+^ cells (D) Quantification of BrdU⁺ NK cell numbers per mg of graft tissue. NKG2D deficiency or blockade and absence of donor Rae-1 decreased NK cell proliferation within the graft. Data indicate the mean ± SD. *P < 0.05 as determined by one-way ANOVA test. (E) ELISA analysis of cytokine levels in graft tissues on day 15 post-transplant. IL-21 levels were increased in AJ grafts to B6.CCR5^⁻/⁻^ recipients compared with the other groups, whereas IL-15 and IL-7 levels showed no significant differences among groups. Data indicate the mean ± SD. *P < 0.05, **P < 0.01 as determined by one-way ANOVA test.

The decreases in NK cell numbers were further reflected by the significant 60-70% decreases in NK cell proliferation within the allografts in the absence of graft Rae-1e or recipient NKG2D (Figure 5C and D). The proliferation of NK cells in B6.CCR5^-/-^ recipients but not in the other allograft-recipient groups on day 15 also correlated with the significantly increased quantities of IL-21 detected in graft homogenates at day 15 post-transplant (Figure 5E). In contrast, quantities of graft IL7 and IL-15 protein were equivalent in allografts from all groups and IL-2 protein was below the level of detection in all allografts (data not shown).

In addition to decreases in proliferation, there were significant decreases in kidney allograft infiltrating NK cell expression of CD107a, an indicator of cytolytic activity, in the absence of either allograft Rae-1e or recipient NKG2D (Figure 6A and B). For those NK cells infiltrating A/J.Rae-1e^-/-^ kidney allografts in B6.CCR5^-/-^ recipients, there was also a decrease in NK cell expression of NKG2D when compared to the NK cells infiltrating wild type A/J kidney allografts (Figure 6C and D).

**Figure 6.**
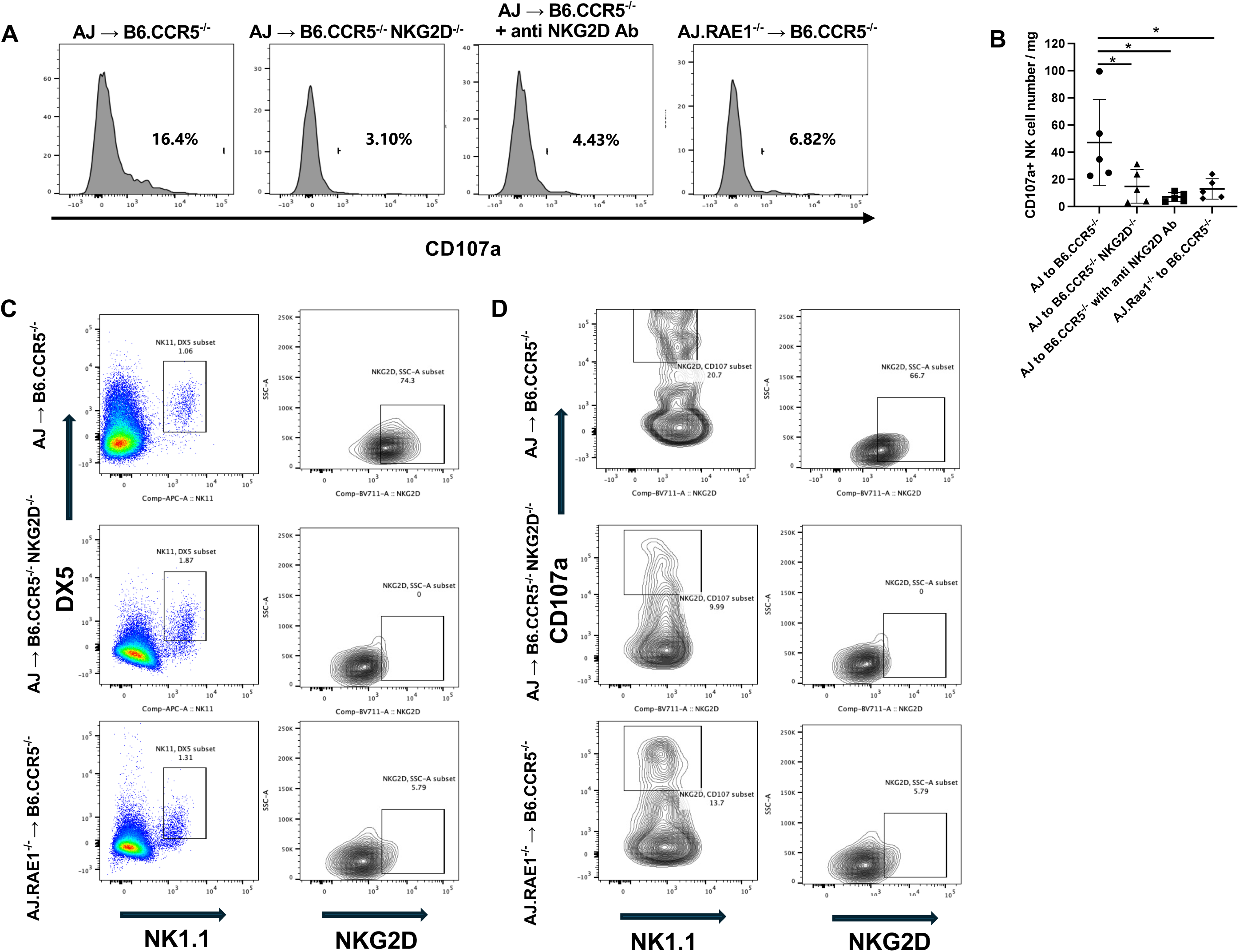
Absence of Rae-1 in donor grafts and recipient NKG2D deficiency attenuate the activation of infiltrating NK cells. (A) Representative histograms of CD107a^+^NK cells in kidney allografts. CD107a expression was assessed within graft-infiltrating NK cells defined as CD45⁺CD3⁻ NK1.1⁺DX5⁺ cells among the groups of AJ grafts to B6.CCR5^/-^, B6.CCR5^⁻/⁻^NKG2D^⁻/⁻^, B6.CCR5^⁻/⁻^ treated with anti-NKG2D mAb recipients and AJ.Rae-1^⁻/⁻^ grafts to B6.CCR5^⁻/⁻^recipients. (B) Quantification of CD107a⁺ NK cell numbers per mg of graft tissue in the groups. NKG2D deficiency or blockade and absence of donor Rae-1 reduced the number of CD107a⁺ NK cell compared with AJ grafts to B6.CCR5^⁻/⁻^. Data indicate the mean ± SD. *P < 0.05, **P < 0.01 as determined by one-way ANOVA test. (C) Representative flow cytometric analysis of NKG2D expression within graft-infiltrating NK cells (CD45⁺CD3⁻NK1.1⁺DX5⁺) and (D) CD107a^+^ NK cell subset in the indicated groups. The expression of NKG2D was reduced in recipient NKG2D-deficient or donor Rae-1-deficient conditions.

In addition to the activation of NK cells in kidney allografts during ABMR in B6.CCR5-/-recipients, our previous reports have documented the coordinate presence of monocytes with an inflammatory (CD11b^hi^Ly6C^hi^) phenotype and the absence of both NK cell activation and inflammatory monocytes in allografts from CCR5^-/-^ recipients with defects in granulocyte and myeloid cell function (11–13). On this basis we tested allografts from each of the recipient groups for the presence of infiltrating inflammatory monocytes on day 15 post-transplant (Figure 7). The presence of these monocytes was clear detectable and significantly higher in wild type A/J kidney allografts from B6.CCR5^-/-^ recipients vs. in A/J allografts from B6.CCR5^-/-^recipients and in A/J.Rae-1e^-/-^ allografts from B6.CCR5^-/-^ recipients. Furthermore, the mean channel fluorescence of Ly6C was significantly higher in the A/J allografts from B6.CCR5^-/-^recipients.

**Figure 7.**
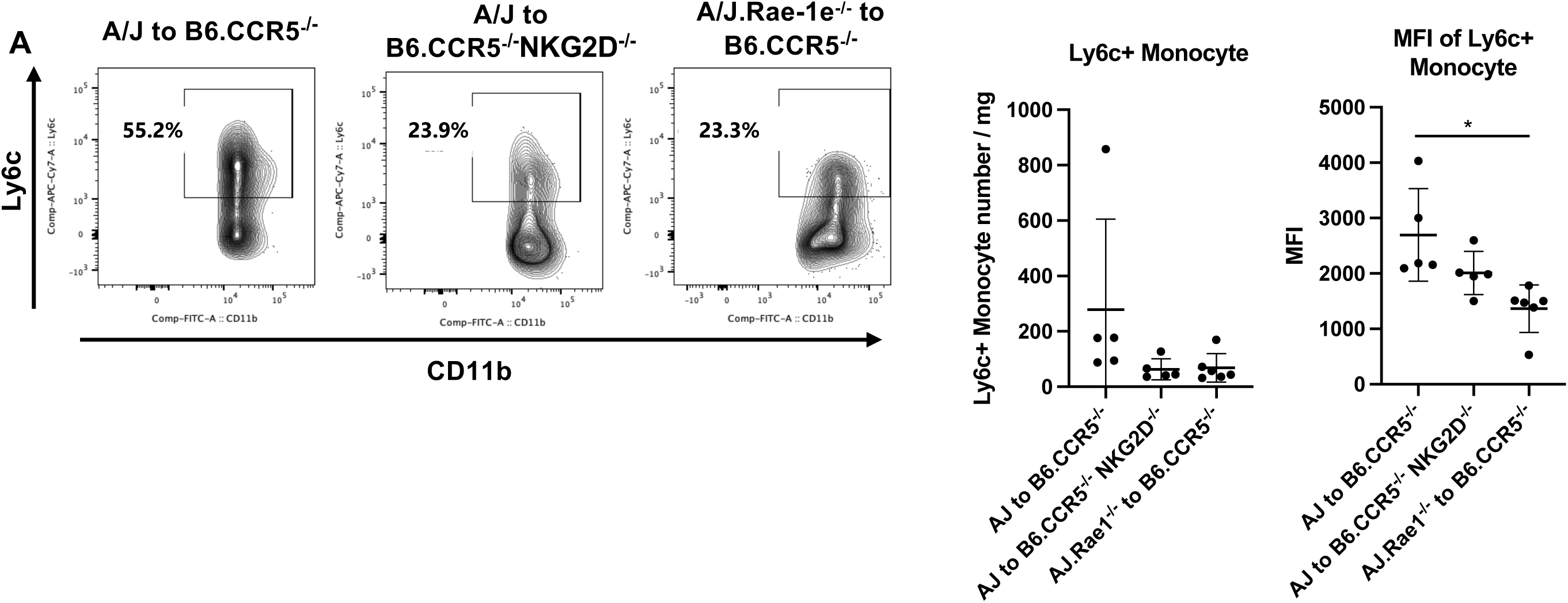
NKG2D signaling is associated with increased accumulation and activation of Ly6C⁺ monocytes in kidney allografts. (A) Representative flow cytometric analysis of graft-infiltrating Ly6C^hi^CD11b^hi^ monocytes. After exclusion of dead cells, leukocytes were gated as CD45⁺ cells, followed by exclusion of CD3⁺ cells. CD11b⁺Ly6G⁻ cells were identified, and F4/80⁻ cells were defined as monocytes. Ly6C expression was evaluated within this monocyte population. Representative contour plots were obtained from AJ grafts to B6.CCR5^/-^, B6.CCR5^⁻/⁻^NKG2D^⁻/⁻^, B6.CCR5^⁻/⁻^ treated with anti-NKG2D mAb recipients and AJ.Rae-1^⁻/⁻^ grafts to B6.CCR5^⁻/⁻^recipients. Quantification of Ly6C⁺ monocyte numbers per mg of graft tissue and Mean fluorescence intensity (MFI) of Ly6C expression in graft-infiltrating monocytes were evaluated in the indicated groups. Rae-1 deficiency grafts reduced MFI of Ly6C expression in monocytes compared with AJ grafts to B6.CCR5^⁻/⁻^. Data indicate the mean ± SD. *P < 0.05 as determined by one-way ANOVA test.

## Discussion

In response to viral infections, mechanisms leading to NK cell activation to express effector functions eliminating virus infected cells is well investigated. Many viruses down-regulate expression of the MHC molecules presenting processed viral peptides and impose intracellular stress that induces expression of ligands for NK activation receptors (20, 21). While investigation into the functions underlying the elimination of virus infected cells continues to provide further insights, fundamental pathways in the process are fairly well delineated. In contrast, clinical evidence points to a clear role for NK cells involvement in mediating acute kidney allograft rejection during ABMR but mechanisms underlying this injury remain unclear and was the focus of this study.

The mouse model used in these studies was the orthotopic transplant of complete MHC mismatched kidneys to CCR5-deficient recipients. In response to the allograft these recipients produce 40-80 fold higher titers of DSA than do wild type recipients of the allografts resulting in aABMR of the allografts by CCR5^-/-^ recipients and not by wild type recipients (8–12). Another relevant feature of this model is that the histopathology of rejection is similar to that observed in clinical kidney grafts experiencing ABMR and the expression of diagnostic transcripts that are associated with the activation of NK cells. In the current report a new insight into this process deals with the allograft expression of the ligand, Rae-1e, required for activation of NK cells during ABMR. We reasoned that expression of this ligand would be induced either during ischemia-reperfusion mediated inflammation in the transplant and sustained to the time of DSA generation or directly by following DSA binding to the vascular endothelium or to other allograft cells. Neither of these scenarios appear to be correct; although Rae-1e expression did not appear in the allograft until serum DSA was detectable, it was not expressed until activation of the innate cellular components mediating the acute graft injury, specifically NK cell and inflammatory monocytes, were apparent. This raises the probability that these innate cells are initially recruited to the allograft and their early activation in the injury process generates the factors inducing Rae-1e expression that, in turn, promotes further and possibly sustained activation of the NK cells within the allograft.

The results also demonstrate the strict requirement for allograft expression of Rae-1e during ABMR and that any expression by allograft infiltrating myeloid or other cells has minimal consequences to the ongoing injury. Furthermore, although NK cells can express multiple activation receptors (20, 22), NK cell activation to mediate aABMR in this kidney transplant model is restricted to expression of NKG2D. It is also worth noting that H60b is a class I MHC-like glycoprotein and also a ligand for NKG2D (23) and was observed at high levels in kidney allografts from all recipient groups in this study but was unable to drive activation of allograft infiltrating NK cells in the absence of Rae-1e. In addition, Rae-1d had the same expression patterns as H60b, but has a lower affinity for NKG2D than Rae-1e and its expression in various tissues may differ with Rae-1e (24). Lin and colleagues (25) have also reported the role of the Rae-1/NKG2D axis in the NK cell mediated development of cardiac allograft vasculopathy in a mouse model where anti-donor class I MHC mAb is administered to recipients of heart allografts and the expression of Rae-1 was detected on graft endothelial cells by flow cytometry (25). Unfortunately, we were unable to stain any kidney allograft cells by flow cytometry or allograft sections for Rae-1 as attempts with several commercially available antibodies did not give detectable or specific staining of sections from allografts expressing Rae-1e transcripts vs from those not expressing the transcripts (e.g. in isografts or Rae-1^-/-^ allografts).

Clinical diagnosis has evolved to utilize detection of transcripts specific for ABMR (14, 15). These transcripts include a set of 6 transcripts associated with the activation of NK cells. We have previously reported that kidney allografts undergoing aABMR in CCR5-deficient recipients also express these transcripts except for granulysin, which is not produced in mice (10, 12). However, the current results indicate the expression of at least two these transcripts in kidney allografts from recipients producing DSA but not able to generate activation of NK cells and inflammatory monocytes within the allograft. The use of these transcripts in the clinic is prompted by graft dysfunction, serum DSA and biopsies indicating the pathology of ABMR and they are clearly not expressed in well-functioning grafts, but the results of this study suggest that DSA on its own may provoke expression in the absence of NK cell and monocyte activation that can go undetected without patient symptoms provoking a biopsy.

We have previously documented the absence of aABMR in allografts transplanted to CCR5^-/-^ recipients with defects in specific myeloid cell functions. Specifically, CCR5^-/-^- MPO^-/-^and -CCR5^-/-^cG^-/-^ recipients do not acutely reject complete MHC mismatched kidney allografts even though the DSA titers in these recipients are equivalent to recipients without these defects (11, 13). There is also a clear absence of NK cell activation as well as the appearance of inflammatory monocytes in the allografts from these myeloid cell defective recipients, suggesting links between the appearance and activation of the two cell populations. Together with the current study, these results suggest that the ABMR process to acute injury and graft failure involves many interacting innate cell components to achieve the function of NK cells and inflammatory monocytes.

Overall, the results of this study indicate the molecular and cellular complexity of aABMR. The expression of Rae-1e to activate NK cells requires more than DSA binding to the vascular endothelium and/or to other allograft cells. Similarly, NK cell activation is accompanied by inflammatory monocyte activation and the absence of one causes the absence of the other. These results suggest initial interactions including DSA induced recruitment of NK cells and monocytes that provoke increases inflammation within the allograft, including Rae-1 expression and/or other activation ligands that mediate the peritubular and glomerular capillary injury leading to graft rejection. Clinical diagnosis is prompted by the injury at a specific time point in the development of the injury and is likely to miss these initial steps in the aABMR process. Further and more extensive studies will need to be performed to accurately delineate the initial states of this allograft injury process.

## Materials and Methods

### Mice

A/J (H-2^a^), C57BL/6 (B6; H-2^b^), B6.CCR5^-/-^ mice and B6.NKG2D^-/-^ mice were obtained from the Jackson Laboratory. B6.Rae-1e^-/-^ mice (26) were obtained from David Raulet, University of California, Berkeley (Berkeley, CA) and a congenic A/J strain carrying the Rae-1e knockout allele was generated by repeated backcrossing to wild type A/J mice for ten generations, selecting Rae-1e^⁺/⁻^ offspring by PCR genotyping at each generation. After the tenth backcross, heterozygous mice were intercrossed to obtain homozygous A/J.Rae-1e^⁻/⁻^ mice, which were maintained by sibling mating. B6.NKG2D^-/-^ mice were crossed with B6.CCR5^-/-^ mice to generate B6.CCR5^-/-^NKG2D^-/-^ mice. Colonies of B6.CCR5^-/-^, B6.CCR5^-/-^NKG2D^-/-^, and A/J.Rae-1e^⁻/⁻^ mice were maintained in the Lerner Research Institute Biological Resources Unit. Male mice (8-12 weeks of age) were used throughout this study.

### Kidney transplantation

Murine orthotopic kidney transplant was performed following the microsurgical methods devised by Zhang and colleagues (27). The donor left kidney was flushed with heparinized Ringer’s solution and harvested *en bloc* with its ureter and vascular pedicle. In the recipient, the native right kidney was removed and the donor artery and vein were anastomosed to the abdominal aorta and inferior vena cava. Ureteral reconstruction of the donor ureter to the recipient bladder was performed as previously described (11, 12). The remaining native left kidney was nephrectomized 4 days after transplantation so that recipient survival was completely dependent on function of the transplanted kidney. Graft survival was assessed by daily observation of recipient health and rejection was confirmed by histopathologic evaluation of the kidney. In some experiments, B6.CCR5^-/-^ recipients of A/J kidney allografts were treated i.p. with 250 µg rat anti-mouse NKG2D monoclonal antibody (catalog BE0351, clone HMG2D, Bio X Cell, NH) on days 8, 13, 18, 23, 30 and 40 after transplantation.

### Immunohistopathology analysis of kidney allografts

Kidney grafts were harvested and fixed in acid methanol (60% methanol and 10% acetic acid). Paraffin-embedded sections (5 μm) were cut and subjected to high temperature antigen retrieval and paraffin removal in Trilogy (Cell Marque-Sigma Aldrich, Rocklin, CA) in a pressure cooker. Endogenous peroxidase activity was eliminated by incubation with 0.03% H_2_O_2_ for 10 minutes, and nonspecific protein interactions were inhibited by incubation with serum-free protein block. The slides were then stained using Periodic acid-Schiff (PAS) or with rabbit polyclonal antiserum to mouse C4d produced as described previously (28). Staining antibodies were visualized using rat or rabbit on mouse HRP-Polymer Kits (Biocare Medical, Pacheco, CA) followed by DAB and counterstained with hematoxylin. Slides were viewed by light microscopy, and images captured using ImagePro Plus (Media Cybernetics, Rockville, MD).

### Flow cytometry analyses of graft infiltrating cells

Harvested kidney grafts were weighed, minced and digested by incubation with type Ⅱ collagenase (catalog 1005021, MP Biomedicals, Santa Ana, CA) and DNase I (catalog 10104159001, MilliporeSigma, Burlington, MA) for 45 minutes at 37°C in RPMI medium. The digested graft was pressed with the plunger of a 3cc syringe, and after 40 μm filtration and centrifugation, the cells were counted and aliquots stained with the following fluorochrome-conjugated antibodies: FITC-anti-CD107a (catalog 121606, clone;1D4B, BioLegend, San Diego, CA), PE-anti-CD49b (catalog 108907, clone; DX5, BioLegend), APC-anti-NK1.1 (catalog 501129655, clone; PK136, eBioscience, San Diego, CA), PE-Cy7-anti-CD3e (catalog 561100, clone; 145-2C11, BD Biosciences, San Jose, CA), BV421-anti-CD45 (catalog 563890, clone; 30-F11, BD Horizon, San Jose, CA), BV711-anti-CD314 (catalog 563694, clone cx5, BD Horizon), FITC-anti-CD11b (catalog 553310, lone;M1/70, BD Biosciences), PE-anti-Ly6G (catalog 551461, clone 1A8, BD Biosciences), APC-anti-CD45 (catalog 559864, clone; 30-F11, BD Biosciences), APC-Cy7-anti-Ly6C (catalog 128026, clone HK1.4, BioLegend), BV711-anti-F4/80 (catalog 123137, clone BM8, BioLegend) and DAPI solution (catalog 564907, BD Biosciences). Cell samples were analyzed on a LSR II or LSR Fortessa X20 flow cytometer (BD Biosciences) and the resulting data analyzed with FlowJo software version 10 (Tree Star Inc., San Carlos, CA). The absolute number of each leukocyte subset was calculated using the following formula: (total number of CD45⁺ leukocytes counted) × (percentage of the target leukocyte population within CD45⁺ cells) / 100. These values are reported as the number of each leukocyte subset per milligram of transplanted tissue. BrdU was administered intraperitoneally (100 µl) on day 14 after transplantation for in vivo labeling and grafts were harvested the following day for flow-cytometric analysis. Intracellular BrdU incorporation was assessed using the FITC BrdU Kit following the manufacturer’s instructions (catalog 559619, BD Biosciences) and flow cytometry of the target cell population.

### Determination of serum DSA titers

Donor-specific IgG antibodies in recipient serum were measured using a flow cytometry–based assay as previously described (8, 9, 29). Briefly, serum was diluted and the different dilutions were incubated with aliquots of kidney graft donor and recipient thymocytes followed by incubation with goat anti-mouse IgG antibody (Jackson ImmunoResearch, West Grove, PA). Antibody-binding was then assessed by flow cytometry. For each serum dilution, the mean channel fluorescence was obtained and the dilution returning the mean channel fluorescence to the level observed when A/J thymocytes were stained with a 1:4 dilution of normal C57BL/6 mouse serum was divided by 2 and reported as the titer.

### RNA extraction, cDNA synthesis, and quantitative analysis of gene expression

Total RNA was isolated from snap-frozen kidney graft tissue using RNeasy Mini Kits (catalog 74004, QIAGEN, Hilden, Germany), and 1 μg was reverse transcribed using High-Capacity cDNA Archive Kits (catalog 4368814, Applied Biosystems, Foster City, CA). Ten microliters of each cDNA was used for quantitative PCR with TaqMan Fast Universal Master Mix and TaqMan primer sets. All PCR reactions were done in duplicate on a 7500 Fast Real-Time PCR System (Applied Biosystems, Thermo Fisher Scientific, Foster City, CA) and quantified using the ddCt method and primers for H60b (Mm04243254_m1), H60c (Mm04243526_m1), IL-33 (Mm00505403_m1), Raet1d (Mm04214213_g1), Raet1e (Mm02530623_s1), IL1RL1 (Mm00516117_m1), IFN-γ (Mm01168134_m1), granzyme B (Mm00442837_m1), perforin (Mm00812512_m1), CCL2 (Mm00441242_m1), CXCR6 (Mm02620517_s1) CX3CR1 (Mm02620111_s1), MyBL1 (Mm00485327_m1) and SH2D1B1 (Mm0468982_ml), with MRPL32 (00777741_SH) as the reference gene (Applied Biosystems, Thermo Fisher Scientific, Waltham, MA).

### Statistical analyses

Data analysis was performed using GraphPad Prism Pro software version10. Comparisons of kidney allograft survival between groups were analyzed using Kaplan-Meier survival curves and log-rank (Mantel Cox) statistics. Statistical differences between 2 experimental groups were analyzed using 2-tailed t tests and comparisons among multiple groups were analyzed by one-way ANOVA. P values less than 0.05 were considered significant. All values for experimental groups are expressed as mean ± SEM.

## Abbreviations

aABMR: acute antibody-mediated rejection
DSA: donor-specific antibody

